# Blinding efficacy and adverse events following repeated transcranial alternating current, direct current, and random noise stimulation

**DOI:** 10.1101/2022.03.04.482999

**Authors:** James G. Sheffield, Sumientra Ramerpresad, Anna-Katharine Brem, Karen Mansfield, Umut Orhan, Michael Dillard, James McKanna, Franziska Plessow, Todd Thompson, Emiliano Santarnecchi, Alvaro Pascual-Leone, Misha Pavel, Santosh Mathan, Roi Cohen Kadosh

## Abstract

As transcranial electrical stimulation (tES) protocols advance, assumptions underlying the technique need to be retested to ensure they still hold. Whilst the safety of stimulation has been demonstrated mainly for a small number of sessions, and small sample size, adverse events (AEs) following multiple sessions remain largely untested. Similarly, whilst blinding procedures are typically assumed to be effective, the effect of multiple stimulation sessions on the efficacy of blinding procedures also remains under question. This is especially relevant in multisite projects where small unintentional variations in protocol could lead to inter-site difference. We report AE and blinding data from 1,019 participants who received up to 11 semi-consecutive sessions of active or sham transcranial alternating current stimulation (tACS), direct current stimulation (tDCS), and random noise stimulation (tRNS), at 4 sites in the UK and US. We found that AEs were often best predicted by factors other than tES, such as testing site or session number. Results from the blinding analysis suggested that blinding was less effective for tDCS and tACS than tRNS. The occurrence of AEs did not appear to be linked to tES despite the use of smaller electrodes or repeated delivery. However, blinding efficacy was impacted in tES conditions with higher cutaneous sensation, highlighting a need for alternative stimulation blinding protocols. This may be increasingly necessary in studies wishing to deliver stimulation with higher intensities.

## 1 Introduction

Transcranial electrical stimulation (tES), the application of weak currents to the scalp to alter neuronal activity (Antal & Herrmann, 2016; Paulus, 2011; Polanía et al., 2018), has been applied to improve human behaviour in typical and atypical populations (Au et al., 2016; Andre R. Brunoni et al., 2017; Grover et al., 2021; Lefaucheur et al., 2017; Nikolin et al., 2015; Salehinejad et al., 2019; Schroeder et al., 2017; Simonsmeier et al., 2018). Despite its increasing application across a range of domains, few studies subject participants to repeated stimulation sessions, potentially due to safety concerns. However, there is substantial potential for it to positively impact cognitive training (Martin et al., 2014; Santarnecchi et al., 2015). Therefore, an effort has been made to confirm that multiple stimulation sessions, mainly using transcranial direct current stimulation (tDCS), lead to limited increases in detrimental or injurious side-effects, known as adverse events (AEs; Nikolin et al., 2017).

It should also be noted that mixed findings within the literature, despite using similar protocols, has generated some scepticism for the technique. Several factors of tES experimental design are often poorly reported, decreasing the reliability of the technique across the literature in general (Filmer et al., 2020). Similarly, in respect to tDCS, a recent study suggests that the blinding procedures currently used may not be as effective as once thought (Turi et al., 2019). Yet, there may be additional variance arising from unexpected sites differences that is not obviously attributable to protocol. Namely, different sites could be using the same protocol but differ in factors such as the experience in running tES experiments, the motivation of the researchers, and the recruited participants.

### 1.1 Adverse events

As with any neuromodulatory technique, questions arise regarding unintended side effects of its application, especially following repeated doses. Until now, the work looking at AEs following tES has been limited, primarily focused on tDCS (Nikolin et al., 2017). Currently, evidence suggest that few, if any, AEs follow when tDCS is given across multiple days (Bikson, Paneri, et al., 2018; Borckardt et al., 2012; Andre Russowsky Brunoni et al., 2011; Matsumoto & Ugawa, 2017). However, as other forms of tES are gaining popularity it is prudent to confirm that both transcranial random noise stimulation (tRNS) and transcranial alternating current stimulation (tACS) are as safe as tDCS, leading to negligible and temporary side-effects. Previous studies suggest that single-session applications of tRNS (Chaieb et al., 2015) are not associated with an increased risk of AEs, whilst tACS is only associated with a minor increase in AEs (Matsumoto & Ugawa, 2017).

Compared to the size of electrodes used most frequently with tES (e.g., 25-35 cm^2^), more manufacturers are moving towards smaller electrode designs (e.g., below 5 cm^2^) to allow for more focal stimulation. Whilst this opens new avenues for research questions, these smaller electrodes lead to greater current densities on the scalp when compared against conventional electrodes (Minhas et al., 2011). Previous research suggests that smaller electrodes are associated with lower reported discomfort than larger electrodes (Fertonani et al 2015), especially with equal current density (Turi et al 2014). However, electrodes reported in these studies are larger than those currently available on the market (9 cm^2^ and 16cm^2^ respectively), especially for those used in joint tES-EEG set-ups. Therefore, as these electrodes are gaining popularity, demonstrating that small surface area electrodes demonstrate similar AE profiles to the larger variants is important.

Moreover, most studies provide tES over a single session. However, potential treatments using tES take several days or weeks. Meta-analysis by Nikolin et al. (2017) suggest that cumulative charge, a product of tDCS intensity, session duration, and number of sessions, did not lead to an increase in AEs. However, this work was limited to tDCS. Similar work has not been conducted for either tRNS or tACS, and as such, it remains an open question as to whether these other tES protocols are equally as safe over repeated administration. It is worth noting that even minor AEs are not only ethically undesirable, but also undermine our ability to successfully blind our participants and experimenters. Therefore, a greater understanding of tES AEs using smaller electrodes and greater number of sessions is vital.

### 1.2 Blinding efficacy

Similar to many interventions, tES depends on effective blinding procedures to ensure changes are due to the stimulation applied rather than participant beliefs regarding stimulation (Fassi & Cohen Kadosh, 2020). Currently, the majority of studies use a fade-in/short-stimulation/fade-out (FSF) method at the beginning of a stimulation session to providing sham stimulation, though the exact parameters of this FSF period vary between studies (Fonteneau et al., 2019). Despite this, some recent studies have demonstrated that FSF may not be as effective at blinding participants as previously shown (Turi et al., 2018).

It is worth noting that some of the factors that may influence the presence of AEs may also impact the blinding efficacy. For example, using smaller electrodes may increase the scalp sensation of stimulation leading to more AEs and worse blinding, though previous work suggests that changes in current density has little impact of cutaneous perception (Fertonani et al., 2015). In contrast. repeated stimulation might lead to fewer active stimulation participants reporting being in the active condition as they become used to the sensation of stimulation.

### 1.3 Multisite studies

Increasingly, multi-site studies are becoming commonplace across a range of domains where intervention studies are the norm. Whilst efforts are often made to ensure homogeneity of practice across sites in these studies, differences may emerge due to variability in expertise or need to adapt to local experimental pressures, amongst other factors. These emergent changes in practice might lead to unintended inter-site variance on measures typically assumed to be stable across sites. For example, in Turi et al. (2018), sites demonstrated differences in their reported impedance values for stimulation. As stimulation devices often attempt to maintain a constant injected current, when impedance values increase, they must increase the voltage. Whilst we would expect higher voltages on the scalp should lead to more sensation (Fertonani et al., 2015), reducing blinding effectiveness, their analysis did not indicate that impedance substantially contributes to stimulation blinding. However, they did note that the model most favourable for predicting blinding included the lab. Whilst this did not demonstrate a clear difference when comparing two labs to their reference lab, there may have been a difference between the two other labs. In contrast, whilst impedance had a near zero effect for predicting participant discomfort during stimulation, site did lead to substantial differences between the reference lab and the two other levels.

Bikson et al.(2018) suggested four reasons for the limited reproducibility in the brain stimulation literature: variation in electrode placement; inconsistencies in electrode preparation; insufficient operator training; insufficient protocol reporting. Assuming that sites develop their protocol together, the last should lead to negligible impact for site differences. However, the other factors may lead to inter-site differences despite the creation of a standardised protocol across all included sites. For example, Bikson et al. (2018) pointed out that small changes to the montage on the scalp (such as those that could occur due to error on the experimenter’s part) can lead to large changes on the cortical surface. Difference in experimenter experience may lead to variance in electrode placement, even when using pre-cut caps to hold electrodes, if these caps are not placed correctly. Differences in skin preparation may affect the electrode impedance and therefore, as explained above, the cutaneous perception of the subject.

Finally, it is important to how experimenter communication may impact stimulation blinding, and Wallace et al. (2016) highlighted that when participants were told that there was an active and sham condition, they were able to guess their stimulation condition at an above chance level (O’Connell et al., 2007), but when they were not told, there was an active and sham condition, participants did not reliably distinguish between conditions (Palmer et al., 2016). Therefore, differences in how the project is discussed with the participant may further influence their perception of the study.

### 1.4 The present study

Here, we report on AEs and blinding data from two large-scale multi-site experiments that involved tES and cognitive training. Participants received either active or sham tDCS, tRNS, or tACS once a day for up to 11 days in a two-week period at one of 4 sites in the UK (University of Oxford[OX]) and the US (Honeywell [HON]; Northeastern University [NEU]; Harvard University [HAR]). Participants were stimulated using electrodes with small surface areas (3.14 cm^2^). These two factors, multiple sessions and small electrodes size might impact participant reports of AEs and blinding. Given the low rate of reported AEs associated with stimulation and the relatively low amplitudes (up to 1.25 mA) at which we delivered stimulation, we did not expect to see a difference between sham and active tES protocols in reported AEs, despite the repeated stimulation. Finally, we investigated differences between sites in both AEs and blinding efficacy with some expectation that these might differ between sites as differences have been observed in previous tDCS experiment (Turi et al., 2019), though we may expect differences between sites in the reported number of AEs and blinding efficacy.

## 2 Methods

### 2.1 Participants

Data were collected from 1,109 participants during a cognitive training study aimed at improving fluid intelligence using adaptive, flexible executive functions training. Participants received between 9-11 tES sessions over a 2-week period at one of four sites: Oxford, UK; Northeastern University, MA, US; Harvard University, MA, US; and Honeywell, MN, US. Participants were excluded from participation in the study if they had a history of migraines, epilepsy, seizure disorders, psychiatric illness, or traumatic brain injury. Participants were asked to refrain from drinking alcohol during the study and avoiding caffeine 1 hour before a testing session. They were also asked to sleep at least 6 hours each night before testing.

### 2.2 Stimulation

Across all conditions, in both experiments, stimulation was delivered by a StarStim8 device (Neuroelectrics, Barcelona, Spain) using circular 3.14 cm^2^ Ag/AgCl electrodes. Stimulation locations were prepared using an abrasive gel (NuPrep) and surgical alcohol, and conductive gel (SignaGel) was used to bridge the gap between the scalp and the electrodes. Impedances were then checked before the testing session began, with it only continuing when impedances were below 10kΩ. In Experiment 1 participants began stimulation on day 3 of the cognitive training, whilst in Experiment 2 stimulation began on day 1. In both experiments, the stimulation condition was pseudonymised to ensure that the experimenters who interacted with the participant during the training were blind to the participants stimulation condition; this information was only know to the experimenters monitoring the data collection at each site. Participants were made aware that they would assigned to either the stimulation or sham conditions during their initial recruitment call and during the first tES-EEG session.

These protocols were selected based on what has been previously successful in the cognitive training literature, as well as settings that have been successful in previous pilot studies across the labs involved.

#### tDCS

1.25 mA tDCS was selected as midpoint between previously successful 1mA (Cohen Kadosh et al., 2010; Floel et al., 2008; Reis et al., 2009) and 1.5mA (Hesse et al., 2007). The tDCS was applied for 30 minutes with the anode placed over F3 and the cathode placed over the AF8 location of the 10-10 electrode system.

#### tRNS

1 mA (peak-to-peak) tRNS (Snowball et al., 2013) was delivered over F3 with the return over F4 for 20 minutes. The tRNS noise was filtered to the 100-500 Hz range.

#### tACS

Two forms of 1 mA (peak-to-peak) theta-frequency (4.5-7 Hz) tACS were delivered over F3 and P3 with a return at Cz. In the first form, tACS was delivered in phase with the endogenous theta rhythm, whilst the tACS was delivered in anti-phase (i.e., 180° difference) with the endogenous theta rhythm in the second form. The frequency of stimulation was adjusted to each individual’s peak theta-band frequency, which was measured using EEG. Both forms of tACS were delivered in 5-minute blocks, with a 10 second break between blocks to estimate theta frequency. In total, participants received 30 minutes of tACS stimulation.

#### Sham

Each active stimulation condition had a corresponding sham condition in which participants received 30 seconds of FSF at the start and end of the stimulation period (Experiment 1), or 20 seconds FSF at the beginning of the stimulation period (Experiment 2) using the same parameters as the active stimulation condition. The tACS sham condition had a random phase with respect to the measured EEG.

**Table 1.**
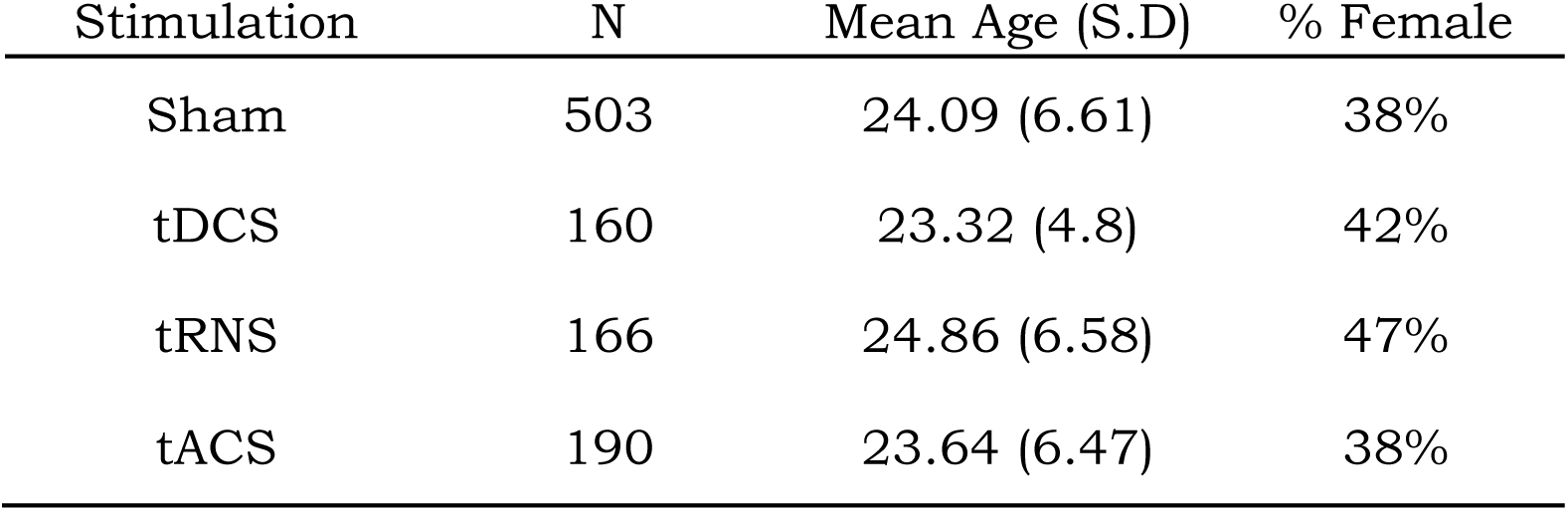
Demographic information for each stimulation group.

### 2.3 Blinding

After participants completed their final session of cognitive training, they were asked to report on whether they believed they received active or sham stimulation (dichotomous response) and the confidence in their response (4-point Likert scale). This can be found in **supplementary materials**.

### 2.4 Adverse Events

At the beginning of each testing session, participants were asked to report on any AEs experienced in the past 24 hours, and at the end of the session, they were asked to report on any AEs experienced during the session. They were asked to report on severity of the AE (4-point Likert scale) and how much they believed it was associated with the stimulation received (7-point Likert Scale). This can be found in **supplementary materials**.

Across both experiments, participants were asked to report on pain, skin irritation, concentration problems, skin sensations (including itching, tingling, and stinging), nervousness, nausea, and any unspecified (other) symptoms. However, there were not enough instances of each AE in all stimulation conditions to run an analysis on them. AEs were only considered for analysis if 3% or more of the sessions within a stimulation group and its associated sham condition resulted in the occurrence of an AE. The AEs that were excluded and the frequencies of each AE are reported in the **supplementary materials**.

### 2.5 Bayesian Inference

We used a Bayesian inference approach because it offers several advantages over frequentist statistics (Wagenmakers et al., 2018). Most relevantly to our study, Bayesian inference is ideal for hypothesis testing as it can quantify evidence towards a particular hypothesis, including a null hypothesis, which frequentist statistics do not. The use of Bayesian inference in brain stimulation studies has been encouraged (Biel & Friedrich, 2018). This is because the results using Bayesian inference allow to find evidence towards the null hypothesis; namely, a lack of AEs and ineffective blinding following stimulation. By comparison, frequentist statistics would only allow us to reject the null hypothesis, but never to accept it. However, the ability to accept the null hypothesis is critical when discussing the adequacy of blinding or the lack of AEs.

### 2.6 Analyses

All analyses were conducted in *R* (R Core Team, 2020) using the *brms* (Bürkner, 2018) and *bayesplot* packages (Gabry et al., 2019).We used multilevel regression analysis with the AE severity or blinding response used as the dependent variable. As most participants reported no AEs, we used models with a zero-inflated negbinomial link function to investigate the effects of stimulation on AE occurrence. We selected priors (normal(0,1)) as we did not have strong expectations regarding a particular effect. Similarly, blinding efficacy was also tested with the *brms* package using Bayesian logistic regression with a Bernoulli link function, and uninformative priors (normal(0,0.5)). As we were interested in a hypothesis testing approach, we calculated the Bayes factor (BF) against an intercept-only null model for each model. The model with the highest BF was selected as long as the BF>3. Based on Jeffreys (2006), BFs>3 were considered as moderate evidence in favour of a model, BFs>10 were considered strong evidence, and BFs>30 were considered extremely strong evidence. Though not of primary interest to this study, elpd_loo-ic_ values are reported in the **supplementary material** for those interested in the out-of-sample predictions of the model.

For our regression models, we defined the following predictors: 1) Site, a categorical variable with 4 levels; Harvard (HAR; the reference level), Oxford (OX), Northeastern (NEU), and Honeywell (HON); 2) Stimulation, a categorical variable with two levels: sham (the reference level), and active stimulation; 3) Session, a continuous variable; and we report the linear change in the effect as a function of session. For each analysis we report the best model vs. the null model, and we report the BF for the model’s predictors. For each predictor in the best model, we provide the regression coefficient (b parameter), with a 95% highest density interval (HDI). For both the AE data and the blinding data, we also provide post-hoc hypothesis tests to break down each interaction effect.

## 3 Results

### 3.1 Adverse Events

#### 3.1.1 tDCS – Experiment 1

The skin sensation AE data following stimulation was most likely under a model containing the main effect for site (BF_10_=3.7^e21^). Both HON (b=1.45, 95%HDI=[0.74, 2.12]) and NEU (b=3.28, 95%HDI=[2.61, 3.92]) demonstrated notably higher sensation AEs following stimulation when compared to HAR. OX demonstrated reduced sensation AEs, but still a higher incidence of sensation AEs compared to HAR (b=0.56, 95%HDI=[-0.24, 1.36]).

For changes in concentration during stimulation, the data was most likely under the model containing the interaction between site and stimulation (BF_10_=3.3^e7^). The interaction terms suggested that HON (b=0.05, 95%HDI=[-1.21, 1.37]) and OX (b=-0.6, 95%HDI=[-2.24, 1.03]) were comparable to HAR in demonstrating fewer concentration problems in the sham group. In contrast, NEU demonstrated substantially higher concentration problems in the sham than the active stimulation group (b=1.13, 95%HDI=[-0.03, 2.28]; **Fig. 1**). Post hoc comparisons confirmed that there was a substantial increase in concentration problems reported by participants in the sham group compared to the active stimulation group at NEU (b=0.68, S.E=0.43, 95%CI=[-0.02,1.39]), which was not seen at HAR (b=-0.45, S.E.=0.52, 95%CI=[-1.30, 0.41]), HON (b=-0.4, S.E.=0.61, 95%CI=[-1.40, 0.59]), or OX (b=-1.05, S.E.=0.89, 95%CI=[-2.55, 0.38]).

**Figure 1:**
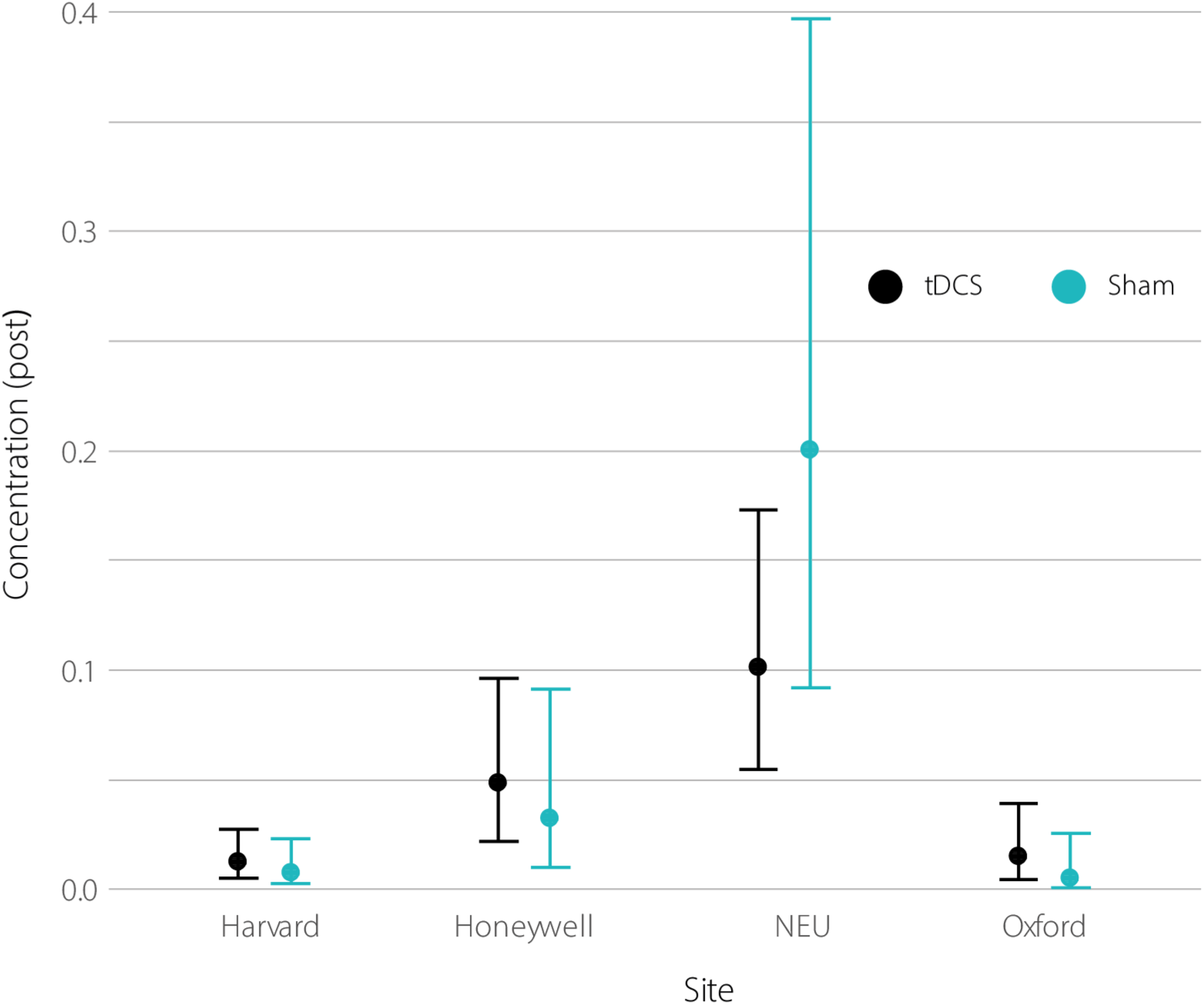
Concentration problems during stimulation as product of active and sham tDCS and site during Experiment 1. Bars denote 95% credible interval for the conditional effect of the site * stimulation effect.

**Figure 2:**
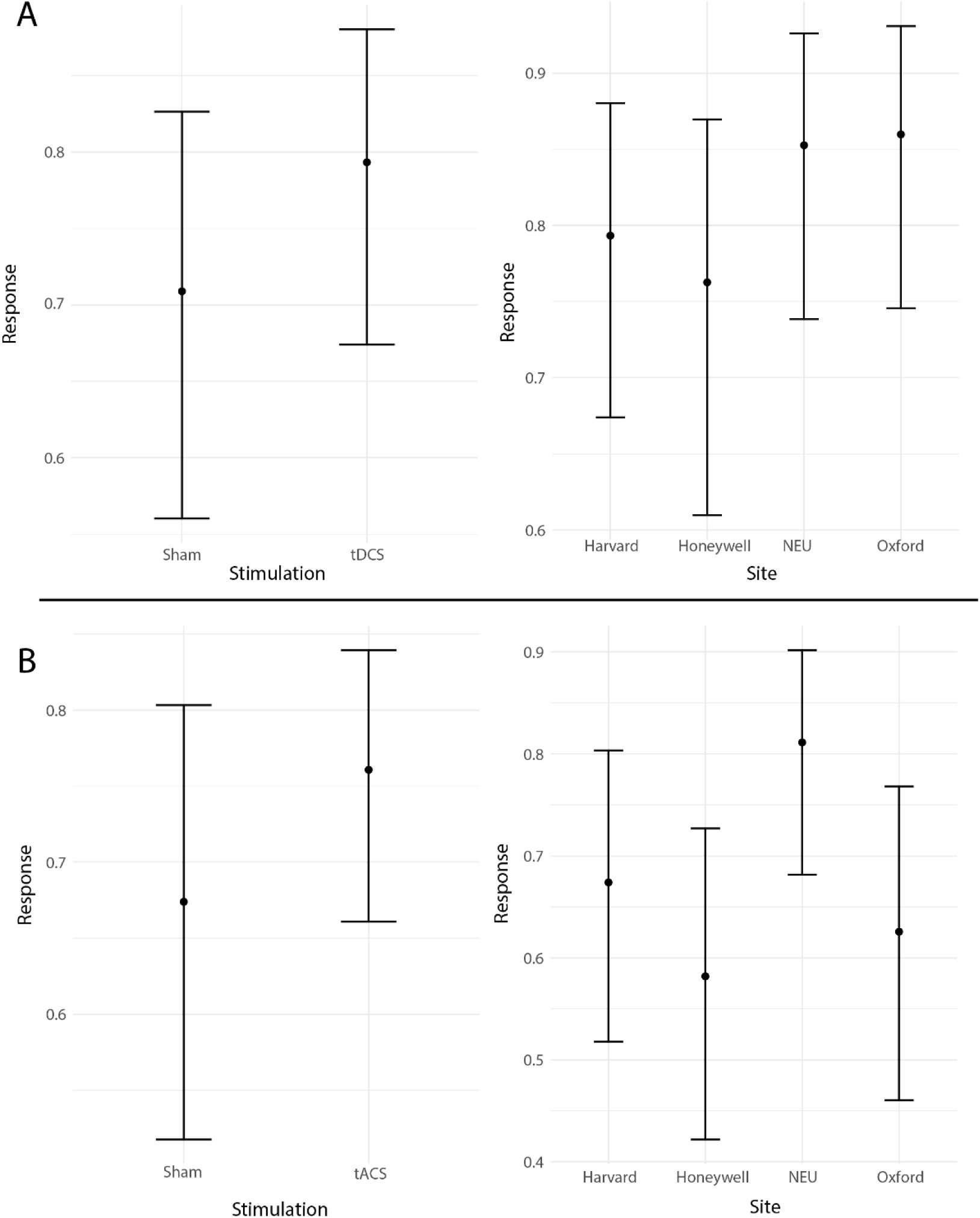
Blinding response demonstrating the main effects for the tDCS and tACS stimulation group. Higher values on the y-axes indicated a tendency to believe receiving active stimulation. A) Site and stimulation main effects for participants from the tDCS group from Experiment 2. B) Site and stimulation main effects for participants from the tACS group (Experiment 2). Bars denote 95% credible interval for the conditional effect of each predictor.

For all other AEs that were entered into the analysis, models containing stimulation were not favourable over the null model or an alternate model not including stimulation (see **supplementary material**). Overall, results from Experiment 1 revealed that the best predictor of participants reports of AEs amongst participants who received either active or sham tDCS was site, with NEU (and HON to a degree) reporting higher rates of AEs than at other sites. However, the concentration AE was best explained by the interaction between stimulation and site, which appeared to be primarily driven by a higher reported number of AEs amongst the sham participants at NEU.

#### 3.1.2 tDCS – Experiment 2

The session alone model was the best for predicting pain (BF_10_=139.23; b=-0.13, 95%HDI=[-.20, -.06]), concentration problems during stimulation (BF_10_=1409.1; b=-0.10, 95%HDI=[-0.14, -0.06]), and unspecified AEs during stimulation (BF_10_=9.12; b=-0.2, 95%HDI=[-0.33, -0.08]). The incidence of all three decreased as sessions increased.

For concentration problems in the 24 hours since the end of the previous session, data indicated that a model containing the main effect of site was the most favourable (BF_10_=1673.17). NEU demonstrated the highest rate of concentration problems in the 24 hours after tDCS session when compared against HAR (b=0.48, 95%HDI=[-0.28, 1.27]), whilst both HON (b=-1.73, 95%HDI=[-2.74, -0.77]) and OX (b=-0.45, 95%HDI=[-1.28, 0.43]) demonstrated lower levels than HAR.

Finally, sensation AE was most likely under the model containing stimulation and session (BF_10_=1.5^e9^). Compared to sham tDCS, active tDCS lead to an increase in sensation AEs (b=0.44, 95%HDI=[0.00, 0.89]), and the number of AEs decreased over sessions (b=-0.12, 95%HDI=[-0.15, -0.08]).

For all other AEs that were entered into the analysis, the null model was favoured over any of the alternative models. Overall, session appeared to be the best predictor of AEs amongst participants in Experiment 2 who received tDCS stimulation, with AE occurrence decreasing over sessions. However, active stimulation did lead to an increase in sensations AEs compared to sham stimulation.

#### 3.1.3 tRNS – Experiment 1

The model that contained the main effect of site was most favourable in the following AEs: 1) pain (BF_10_=521.44): Both HON (b=1.68, 95%HDI=[0.68, 2.69]) and NEU (b=1.40, 95%HDI=[0.32, 2.42]) reported substantially higher AE incidence than HAR. By comparison, OX demonstrated fewer pain AEs (b=-0.80, 95%HDI=[-2.31, 0.57]). 2) Concentration problems during tRNS (BF_10_=6.3^e6^): both HON (b=1.60, 95%HDI=[0.68, 2.60]) and NEU (b=2.23, 95%HDI=[1.35, 3.13]) reported substantially higher rates of concentration AEs compared to HAR, whilst OX demonstrated fewer (b=-0.91, 95%HDI=[-2.34, 0.46]). 3) Sensation (BF_10_=3.7^e31^): both HON (b=1.73, 95%HDI=[0.96, 2.41]) and NEU (b=3.33, 95%HDI=[2.67, 4]) reported substantially higher reported incidence of sensations compared to HAR. OX reported substantially fewer sensation AEs (b=-1.33, 95%HDI=[-2.67, -0.07]).

Finally, for unspecified AEs the model with both the site and stimulation main effect were most favourable (BF_10_=913.97). Participants in the active stimulation condition were more likely to report AEs than those in the sham condition (b=0.99, 95%HDI=[0.13, 1.86]). NEU demonstrated the highest incidence of unspecified AEs (b=1.28, 95%HDI=[0.37, 2.21]), with HON (b=0.91, 95%HDI=[-0.10, 1.87]) also demonstrating higher rates than HAR. OX reported the lowest rates of AEs (b=-0.99, 95%HDI=[-2.41, 0.35]).

For all other AEs that were entered into the analysis, the null model was favoured over the alternate models. Similar to the tDCS Experiment 1 findings, the findings from tRNS participants in Experiment 1 suggest that the best predictors of AE occurrence was site, with the HON and NEU sites demonstrating higher incidence of AEs than the baseline HAR group. However, unspecified AEs were higher amongst participants who received active tRNS stimulation compared to sham stimulation at some sites.

#### 3.1.4 tRNS – Experiment 2

The session alone model was the best for predicting pain during tRNS (BF_10_=232391.50; b=-0.17, 95%HDI=[-0.23, -0.11]), sensation during tRNS (BF_10_=208300.87 ;b=-0.12, 95%HDI=[-0.16, -0.08]), and concentration during tRNS (BF_10_=23409.78; b=-0.09, 95%HDI=[-0.13, -0.06]). Similar to the tDCS Experiment 2 findings, AE incidence decreased with the number of sessions.

By comparison, concentration problems in the 24 hours after tRNS were most likely under a model containing site (BF_10_=3173.43). Relative to HAR, higher rates of concentration AEs were likely at NEU (b=0.56, 95%HDI=[-0.27, 1.46]), whilst lower rates were likely at HON (b=-1.54, 95%HDI=[-2.55, -0.53]) and OX (b=-1.13, 95%HDI=[-2.13, -0.07]).

For all other AE types that were entered into the analysis, the null model was favoured over the alternate models. Much like tDCS participants in Experiment 2, session appeared to be the best predictor of AEs amongst participants who received active or sham tRNS in Experiment 2, with a decrease in AE occurrence over sessions.

#### 3.1.5 tACS

tACS was run only in Experiment 2. Similar to the other two forms of stimulation run in Experiment 2, session alone appeared to be the best predictor of AE, in this case for the occurrence for scalp irritation in the 24 hours period after tACS (BF_10_=1.9^e10^; b=-0.2, 95%HDI=[-0.26, -0.15]), sensations during tACS (BF_10_=7.2^e12^; b=-0.13, 95%HDI=[-0.16, -0.10]), and unspecified AEs experienced (BF_10_=270778.75; b=-0.31, 95%HDI=[-0.42, -0.20]), with increasing session number leading to a decrease in AE occurrence.

For all other AE types that were entered into the analysis, the null model was favoured over the alternate models or there was no clear evidence towards one or the other. Overall, as with the other two Experiment 2 stimulation conditions, session was the best predictor of AEs with participants reporting decreases in AEs as the sessions.

### 3.2 Stimulation blinding

#### 3.2.1 tDCS- Experiment 1

The data was equally likely under the stim only model compared to the null model (BF_10_ = 1.48). However, the null model was favourable all other alternate models (site only: BF_10_ = 0.07 site and stim: BF_10_ = 0.09; site by stim: BF_10_ = 0.06).

#### 3.2.2 tDCS-Experiment 2

The data were most likely under the model containing the main effects of site and stimulation (BF_10_ = 3.03). Stimulation was more likely to lead to active reporting (b = 0.45, 95%HDI = [-0.20, 1.06]). Whilst HON was comparable to HAR (b = -0.18, 95%HDI = [-0.89, 0.55]), both NEU (b = 0.42, 95%HDI = [-0.31, 1.17]) and OX (b = 0.48, 95%HDI = [-0.25, 1.22]) were more likely to report receiving active stimulation across both stimulation conditions.

#### 3.2.3 tRNS-Experiment 1

The data was equally likely under any of the alternate models as the null model (stim only: BF_10_ = 0.63; site only: BF_10_ = 0.71; site and stim: BF_10_ = 0.44; site by stim: BF_10_ = 0.39)

#### 3.2.4 tRNS-Experiment 2

The data were moderately more likely under the null model than the model containing the stim by site interaction (BF_10_ = 0.21), likely due to increased complexity of the interaction term. However, the likelihood of the null model was only anecdotal under the model containing both the main effects (BF_10_ = 0.39), the site effect alone (BF_10_ = 0.44), or the stimulation effect alone (BF_10_ = 0.90).

#### 3.2.5 tACS

For tACS, the data was most likely under a model with the main effects of site and stimulation (BF10 = 60.07). Stimulation led to a slight increase in participants reporting that they were in the active stimulation condition (b = 0.43, 95%HDI = [-0.15, 1.03]). In comparison to participants at HAR, participants at NEU were more likely to report being in the active stimulation condition (b = 0.74, 95%HDI = [0.09, 1.38]). Participants at OX (b = -0.21, 95%HDI = [-0.84,0.41]) and HON (b = -0.40, 95%HDI = [-1.01, 0.20]) were largely comparable to HAR participants.

## 4 Discussion

In this study we examined the effects of tDCS, tRNS, and tACS on blinding and AEs at 4 different sites and on 1,019 participants in two multi-session tES experiments. Our AE findings are in line with previous studies that suggest that the likelihood of AEs following a mild level tES are low (Chaieb, Antal, & Paulus, 2015; Matsumoto & Ugawa, 2017). We have demonstrated that this remains true following repeated tES dosages using small surface area (and thus higher current density) electrodes, and this is further reinforced by the relatively low incidence of some AEs across the entire experiment (<1% incidence across all sessions; see **supplementary materials**). This is in line with previous research that support the safety of smaller electrodes for tES (Fertonani et al., 2015; Turi et al, 2014). Whilst tES led to an increased likelihood of a few AEs, both site and session across tES conditions, appeared to be a more consistent pair of predictors of AE likelihood. Specifically, some sites had higher AEs than others, and AE incidence decreased over the study period.

Generally, stimulation was not associated with increased occurrence of AEs. However, there were certain AEs (e.g. sensation and concentration AEs) that were increased following active stimulation compared to sham. Though it must be noted that the occurrence of any of these AEs were low when considering the total number of sessions under consideration (see **supplementary tables 2-6**). Across 7,932 sessions, 3,833 individual AE were reported (48% of sessions, assuming participants only reported 1 AE each session). Participant reports of skin sensations were by far the most common AE (approximately 40% of all reported AEs), which is to be expected considering that most participants report perceptual response to stimulation.

Notably, we saw a division between the two experiments in the best predictor of stimulation AEs. In Experiment 1 site was the most reliable predictor for AE incidence, whilst in Experiment 2 session was the best predictor. One potential explanation, as highlighted by Bikson et al. (2018), is that experimenter experience may be one of the factors that lead to substantial site differences in Experiment 1. Both the HON and NEU sites, who had the lowest levels of experience with stimulation at the start, often had participants reporting the highest levels of AEs. This may have led to a higher number of AEs as the lower level of experience may have led to worse participant preparation and therefore higher impedance values at the scalp, resulting in more AEs. Potentially, lower expertise may lead to higher reporting of AEs as the experimenter is more cautious about any potentially AEs the participant mentions, even if they are unlikely to be associated with the stimulation itself. Alternatively, subtle differences in the way questions regarding AEs were presented to participants may have led to higher numbers of reported AEs (Wallace et al., 2016). The former is unlikely, as all sites used the same stimulation software, which requires the electrode impedance to be below a specified limit to allow starting the stimulation. Anecdotally, experimenters at NEU elaborated on AE questions more than those at the other sites. As such, these findings may highlight the importance of standardised AE reporting tools and protocols across studies. This is especially relevant here, as despite using similar protocols between sites (including AE reporting protocols), we found notable differences. Whilst all sites involved in the study collaborated in the design and implementation of the protocols, and met biweekly for phone calls to exchange knowledge, experimenters did not travel between sites, and this may be key for successful implementation of multisite methodologies.

In contrast to this, the best predictor of AEs in Experiment 2 was the number of sessions of stimulation the participant had received. The occurrence of AEs was higher in the earlier sessions compared to the later sessions, which is partially in-line with Nikolin et al. (2017), who found that cumulative tDCS charge was not associated with AE increases. The higher rate of AEs in earlier sessions may be due to habituation to stimulation leading to a reduced skin response in later sessions. AEs they may have originally reported in early sessions might seem less intrusive in later sessions, and therefore may be ignored. In contrast to Experiment 1, participants in Experiment 2 received stimulation from the first session rather than day 3. However, whether these additional two days of stimulation is enough to increase such a habituation effect is unclear.

We found that stimulation blinding was ineffective for tACS over both experiments and for tDCS during the second experiment. Similarly, whilst participant confidence in their response generally increased in the active stimulation condition compared to the sham condition, there was notable variation between sites in each condition (see **supplementary materials**). For tACS and tDCS, models that included both stimulation and experimental site were preferred models over the null, with participants in the active stimulation condition being more likely to report being in the active condition than their sham counterparts. The perceptual threshold for tDCS, in some participants, has been previously demonstrated to be around 0.4 mA (Ambrus et al., 2010), and anecdotally, we have noted that participants are most sensitive to tACS compared to other forms of tES. As such, it is unsurprising that the blinding was least effective for these conditions.

By comparison, our data provided no clear evidence for either the null or alternate models for tRNS in either experiments, or for tDCS in the first experiment, suggesting that our study was not sensitive enough to detect differences (or lack thereof) between the conditions. Potentially, the lack of clear stimulation effect on tRNS blinding may be due to the lower amplitude (compared to tDCS, where the stimulation current was approximtaly twice as large) or shorter duration of stimulation (compared to tDCS/tACS). However, instead of equating the stimulation protocols in terms of current strength and duration, we chose the parameters that were most likely to be effective to lead to cognitive effects in our cognitive training project. Therefore, our conclusions do not indicate that tRNS is “better” than tDCS or tACS in terms of blinding, but with the set of the parameters that we used, tRNS is less likely to lead to ineffective blinding, while tDCS and tACS using the parameters that we used here could compromise blinding. It should be noted that, in comparison to previous stimulation studies that examined blinding, our study contained a larger sample size. With Bayesian statistics, larger sample sizes create a more accurate representation of the credible intervals and a better approximation of the of the posteriors (De Santis & Gubbiotti, 2021). The larger sample size reported here should lead to a more accurate approximation of the impact of stimulation on blinding.

These results suggest that FSF blinding protocols are less effective than initially thought for tACS, and potentially for tDCS, and so alternate methods must be sought. One proposed solution is to use local anaesthetics or other pharmacological interventions to aid in participant blinding, and these have been used to ameliorate the cutaneous sensation of stimulation previously (Guleyupoglu et al., 2014; Jamil et al., 2017), though may not be entirely successful for high-current stimulation (Samani, Agboada, Jamil, Kuo & Nitsche, 2019). However, participant response of uncomfortable skin sensations may be one of the early indicators of potential AEs. Though erythema, and even rarer, blistering is uncommon, participants alerting experimenters to their discomfort leading to early termination of stimulation may be one of the factors in their rarity. Moreover, this may be ineffectual in situations where stimulation blinding is due to other perceptual side-effects, such as phosphenes or auditory nerve activation cause by tACS (Zeng, Tran, Richardson, Sun, and Xu, 2019). Our view is that from an ethical standpoint it may not be a reasonable answer to risk participant health for successful blinding. Therefore, it is on us of the stimulation community to come up with a scientifically and ethically viable alternative. That said, there is some evidence suggesting that different analgesic approaches may be more appropriate than others for preventing AEs, at least for tDCS (Guarienti et al., 2015) and in a review of the safety and ethical concerns of stimulation, Antal et al. (2017) highlight that pharmacological interventions have been used with stimulation to no adverse effect. At the same time, such solution could lead to a scenario that most, if not all, the participants will believe that they do not receive stimulation. It is an open question on how such procedure would impact the results, and potential translation, given that patients who will receive tES as part of a future treatment will know that they receive active form of stimulation. Overall, these results suggest that within certain stimulation conditions, blinding may be ineffective, and these failures to appropriately blind may be a driving factor behind the largely variable tES efficacy in the literature.

One potential limitation of our study is that the differences between the different sites may be due to different populations at each site. Even two of the sites (NEU and HAR) that recruited from the same geographical location (Boston area) demonstrated large differences in the demographics of their participants (see **supplementary table 7**). However, even if such a limitation is true, it is essential to consider how such site differences would be minimised to reduce the impact of ineffective blinding in order to improve basic and clinical studies. Alternatively, these differences may have been due to variation in the way the experimenters discussed the study with the participants (Wallace et al., 2016). As mentioned above, anecdotally, investigators at NEU provided more detail than investigators at other sites when answering questions. These differences in information may account for some of the inter-site variance, highlighting the need to standardised protocols, including answers to frequently asked questions, across all sites in a study.

A second important limitation is that the stimulation conditions differed in their montage, stimulation intensity, and duration as these were the stimulation profiles thought most likely to affect significant change in the main experiment (fluid intelligence change during cognitive training). However, this renders direct comparison between the stimulation conditions impossible.

Finally, whilst participants were offered to leave a reason for why they believed they were in a specific stimulation condition, this was not enforced, and their answers were freeform. As such, we cannot draw conclusions regarding why blinding failed for tACS and tDCS here. Further work using a structured stimulation blinding debrief questionnaire may help elucidate the reasons for blinding failure leading to improved techniques to ensure blinding.

### 4.1 Conclusions

In summary, whilst our results provide evidence to support the safety of chronic tES protocols using small surface area electrodes, they suggested an issue with commonly accepted blinding protocols for tDCS and tACS. These inconsistencies in effective blinding noted across the two experiments may help to understand the variance in the tDCS literature. As such, a need for alternative blinding methods better suited for tES protocols with higher cutaneous sensation may be necessary. Issues with multisite blinding differences were unexpected but highlight additional consideration for further research. Differences in experimenter-participant communication may have led to differences in AE reporting between sites, despite the use of a standardised protocol. If these issues might arise between sites with extensive communication to maintain parity, then it raises greater concern about the potential for comparison between studies without direct communication and comparison during the protocol creation and data collection phases. Therefore, methods for compensating for this variance between studies need to be developed.

## Acknowledgements

We would like to thank Giulia Maistrello, Margherita Nulli, Francesco Sella, Thomas Reed, and Morio Hamada (Oxford); Dennis Cornhill, Sandi White, Zane Thimmesch-Gill, Jessamy Almquist, and Forrest Olsen (Honeywell); Erica Levenbaum, Alexandra Emmendorfer and Ann Connor (Harvard); Asieh Ahani, Deniz Erdogmus, Sadegh Salehi, Yeganeh Marghi, and Seyhmus Güler (Northeastern University); Eben Myers, Garrett Kimball, Adam Chizmar, Michael Wagner, and Jessica Trybus (Simcoach Games) for their valuable help. The WIN is supported by core funding from the Wellcome Trust (203139/Z/16/Z). This research is based upon work supported by the Office of the Director of National Intelligence (ODNI), Intelligence Advanced Research Projects Activity(IARPA), via 2014-13121700007. The views and conclusions contained herein are those of the authors and should not be interpreted as necessarily representing the official policies or endorsements, either expressed or implied, of the ODNI, IARPA, or the U.S. Government. The U.S. Government is authorised to reproduce and distribute reprints for Government purposes notwithstanding any copyright annotation thereon.

## Reference

Ambrus, G. G., Paulus, W., & Antal, A. (2010). Cutaneous perception thresholds of electrical stimulation methods: Comparison of tDCS and tRNS. Clinical Neurophysiology, 121(11), 1908–1914. https://doi.org/10.1016/j.clinph.2010.04.020

Antal, A., & Herrmann, C. S. (2016). Transcranial Alternating Current and Random Noise Stimulation: Possible Mechanisms. Neural Plasticity, 2016. https://doi.org/10.1155/2016/3616807

Au, J., Katz, B., Buschkuehl, M., Bunarjo, K., Senger, T., Zabel, C., Jaeggi, S. M., & Jonides, J. (2016). Enhancing Working Memory Training with Transcranial Direct Current Stimulation Jacky. Journal of Cognitive Neuroscience, 28(9), 1419–1432. https://doi.org/10.1162/jocn

Biel, A. L., & Friedrich, E. V. C. (2018). Why you should report bayes factors in your transcranial brain stimulation studies. Frontiers in Psychology, 9(JUL), 1–4. https://doi.org/10.3389/fpsyg.2018.01125

Bikson, M., Brunoni, A. R., Charvet, L. E., Clark, V. P., Cohen, L. G., Deng, Z. De, Dmochowski, J., Edwards, D. J., Frohlich, F., Kappenman, E. S., Lim, K. O., Loo, C., Mantovani, A., McMullen, D. P., Parra, L. C., Pearson, M., Richardson, J. D., … Lisanby, S. H. (2018). Rigor and reproducibility in research with transcranial electrical stimulation: An NIMH-sponsored workshop. Brain Stimulation, 11(3), 465–480. https://doi.org/10.1016/j.brs.2017.12.008

Bikson, M., Paneri, B., Mourdoukoutas, A., Esmaeilpour, Z., Badran, B. W., Azzam, R., Adair, D., Datta, A., Fang, X. H., Wingeier, B., Chao, D., Alonso-Alonso, M., Lee, K., Knotkova, H., Woods, A. J., Hagedorn, D., Jeffery, D., … Tyler, W. J. (2018). Limited output transcranial electrical stimulation (LOTES-2017): Engineering principles, regulatory statutes, and industry standards for wellness, over-the-counter, or prescription devices with low risk. Brain Stimulation, 11(1), 134–157. https://doi.org/10.1016/j.brs.2017.10.012

Borckardt, J. J., Bikson, M., Frohman, H., Reeves, S. T., Datta, A., Bansal, V., Madan, A., Barth, K., & George, M. S. (2012). A pilot study of the tolerability and effects of high-definition transcranial direct current stimulation (HD-tDCS) on pain perception. Journal of Pain, 13(2), 112–120. https://doi.org/10.1016/j.jpain.2011.07.001

Brunoni, Andre R., Moffa, A. H., Sampaio-Junior, B., Borrione, L., Moreno, M. L., Fernandes, R. A., Veronezi, B. P., Nogueira, B. S., Aparicio, L. V. M., Razza, L. B., Chamorro, R., Tort, L. C., Fraguas, R., Lotufo, P. A., Gattaz, W. F., Fregni, F., & Benseñor, I. M. (2017). Trial of Electrical Direct-Current Therapy versus Escitalopram for Depression. New England Journal of Medicine, 376(26), 2523–2533. https://doi.org/10.1056/nejmoa1612999

Brunoni, Andre Russowsky, Amadera, J., Berbel, B., Volz, M. S., Rizzerio, B. G., & Fregni, F. (2011). A systematic review on reporting and assessment of adverse effects associated with transcranial direct current stimulation. International Journal of Neuropsychopharmacology, 14(8), 1133–1145. https://doi.org/10.1017/S1461145710001690

Bürkner, P. C. (2018). Advanced Bayesian multilevel modeling with the R package brms. R Journal, 10(1), 395–411. https://doi.org/10.32614/RJ-2018-017

Chaieb, L., Antal, A., & Paulus, W. (2015). Transcranial random noise stimulation-induced plasticity is NMDA-receptor independent but sodium-channel blocker and benzodiazepines sensitive. Frontiers in Neuroscience, 9(APR), 1–9. https://doi.org/10.3389/fnins.2015.00125

Cohen Kadosh, R., Soskic, S., Iuculano, T., Kanai, R., & Walsh, V. (2010). Modulating neuronal activity produces specific and long-lasting changes in numerical competence. Current Biology, 20(22), 2016–2020.

De Santis, F., & Gubbiotti, S. (2021). Sample Size Requirements for Calibrated Approximate Credible Intervals for Proportions in Clinical Trials. International journal of environmental research and public health, 18(2), 595. https://doi.org/10.3390/ijerph18020595

Fassi, L., & Kadosh, R. C. (2020). Is it all in our head? When subjective beliefs about receiving an intervention are better predictors of experimental results than the intervention itself. BioRxiv.

Fertonani, A., Ferrari, C., & Miniussi, C. (2015). What do you feel if I apply transcranial electric stimulation? Safety, sensations and secondary induced effects. Clinical Neurophysiology, 126(11), 2181–2188.

Filmer, H. L., Mattingley, J. B., & Dux, P. E. (2020). Modulating brain activity and behaviour with tDCS: Rumours of its death have been greatly exaggerated. Cortex, 123, 141–151. https://doi.org/10.1016/j.cortex.2019.10.006

Flöel, A., Rösser, N., Michka, O., Knecht, S., & Breitenstein, C. (2008). Noninvasive brain stimulation improves language learning. Journal of cognitive neuroscience, 20(8), 1415–1422.

Fonteneau, C., Mondino, M., Arns, M., Baeken, C., Bikson, M., Brunoni, A. R., Burke, M. J., Neuvonen, T., Padberg, F., Pascual-Leone, A., Poulet, E., Ruffini, G., Santarnecchi, E., Sauvaget, A., Schellhorn, K., Suaud-Chagny, M. F., Palm, U., & Brunelin, J. (2019). Sham tDCS: A hidden source of variability? Reflections for further blinded, controlled trials. Brain Stimulation, 12(3), 668–673. https://doi.org/10.1016/j.brs.2018.12.977

Gabry, J., Simpson, D., Vehtari, A., Betancourt, M., & Gelman, A. (2019). Visualization in Bayesian workflow. Journal of the Royal Statistical Society. Series A: Statistics in Society, 182(2), 389–402. https://doi.org/10.1111/rssa.12378

Grover, S., Nguyen, J. A., Viswanathan, V., & Reinhart, R. M. G. (2021). High-frequency neuromodulation improves obsessive–compulsive behavior. Nature Medicine, 27(2), 232–238. https://doi.org/10.1038/s41591-020-01173-w

Guarienti, F., Caumo, W., Shiozawa, P., Cordeiro, Q., Boggio, P. S., Benseñor, I. M., Lotufo, P. A., Bikson, M., & Brunoni, A. R. (2015). Reducing Transcranial Direct Current Stimulation-Induced Erythema With Skin Pretreatment: Considerations for Sham-Controlled Clinical Trials. Neuromodulation, 18(4), 261–265. https://doi.org/10.1111/ner.12230

Guleyupoglu, B., Febles, N., Minhas, P., Hahn, C., & Bikson, M. (2014). Reduced discomfort during high-definition transcutaneous stimulation using 6% benzocaine. Frontiers in neuroengineering, 7, 28.

Hesse, S., Waldner, A., Mehrholz, J., Tomelleri, C., Pohl, M., & Werner, C. (2011). Combined transcranial direct current stimulation and robot-assisted arm training in subacute stroke patients: an exploratory, randomized multicenter trial. Neurorehabilitation and neural repair, 25(9), 838–846.

Jamil, A., Batsikadze, G., Kuo, H. I., Labruna, L., Hasan, A., Paulus, W., & Nitsche, M. A. (2017). Systematic evaluation of the impact of stimulation intensity on neuroplastic after-effects induced by transcranial direct current stimulation. The Journal of physiology, 595(4), 1273–1288.

Jeffreys, H. (2006). Theory of probability. In Community Care (Issue 1638, pp. 26–27). Clarendon Press. https://global.oup.com/academic/product/the-theory-of-probability-9780198503682?cc=gb&lang=en&

Lefaucheur, J. P., Antal, A., Ayache, S. S., Benninger, D. H., Brunelin, J., Cogiamanian, F., Cotelli, M., De Ridder, D., Ferrucci, R., Langguth, B., Marangolo, P., Mylius, V., Nitsche, M. A., Padberg, F., Palm, U., Poulet, E., Priori, A., … Paulus, W. (2017). Evidence-based guidelines on the therapeutic use of transcranial direct current stimulation (tDCS). Clinical Neurophysiology, 128(1), 56–92. https://doi.org/10.1016/j.clinph.2016.10.087

Martin, D. M., Liu, R., Alonzo, A., Green, M., & Loo, C. K. (2014). Use of transcranial direct current stimulation (tDCS) to enhance cognitive training: effect of timing of stimulation. Experimental Brain Research, 232(10), 3345–3351. https://doi.org/10.1007/s00221-014-4022-x

Matsumoto, H., & Ugawa, Y. (2017). Adverse events of tDCS and tACS: A review. Clinical Neurophysiology Practice, 2, 19–25. https://doi.org/10.1016/j.cnp.2016.12.003

Minhas, P., Datta, A., & Bikson, M. (2011). Cutaneous perception during tDCS.pdf. Clinical Neurophysiology, 122(4), 637–638. https://doi.org/10.1016/j.clinph.2010.09.023.Cutaneous

Nikolin, S., Huggins, C., Martin, D., Alonzo, A., & Loo, C. K. (2017). Safety of repeated sessions of transcranial direct current stimulation: A systematic review. Brain Stimulation, 17, 278–288. https://doi.org/10.1016/j.brs.2017.10.020

Nikolin, S., Loo, C. K., Bai, S., Dokos, S., & Martin, D. M. (2015). Focalised stimulation using high definition transcranial direct current stimulation (HD-tDCS) to investigate declarative verbal learning and memory functioning. NeuroImage, 117, 11–19. https://doi.org/10.1016/j.neuroimage.2015.05.019

O’Connell, R. G., Dockree, P. M., Bellgrove, M. A., Kelly, S. P., Hester, R., Garavan, H., Robertson, I. H., & Foxe, J. J. (2007). The role of cingulate cortex in the detection of errors with and without awareness: A high-density electrical mapping study. European Journal of Neuroscience, 25(8), 2571–2579. https://doi.org/10.1111/j.1460-9568.2007.05477.x

Palmer, C., Zapparoli, L., & Kilner, J. M. (2016). A New Framework to Explain Sensorimotor Beta Oscillations. Trends in Cognitive Sciences, 20(5), 321–323. https://doi.org/10.1016/j.tics.2016.03.007

Paulus, W. (2011). Transcranial electrical stimulation (tES - tDCS; tRNS, tACS) methods. Neuropsychological Rehabilitation, 21(5), 602–617. https://doi.org/10.1080/09602011.2011.557292

Polanía, R., Nitsche, M. A., & Ruff, C. C. (2018). Studying and modifying brain function with non-invasive brain stimulation. Nature Neuroscience, 21(2), 174–187. https://doi.org/10.1038/s41593-017-0054-4

R Core Team. (2020). R: A Language and Environment for Statistical Computing. https://www.r-project.org

Reis, J., Schambra, H. M., Cohen, L. G., Buch, E. R., Fritsch, B., Zarahn, E., … & Krakauer, J. W. (2009). Noninvasive cortical stimulation enhances motor skill acquisition over multiple days through an effect on consolidation. Proceedings of the National Academy of Sciences, 106(5), 1590–1595.

Salehinejad, M. A., Wischnewski, M., Nejati, V., Vicario, C. M., & Nitsche, M. A. (2019). Transcranial direct current stimulation in attention-deficit hyperactivity disorder: A meta-analysis of neuropsychological deficits. PLOS ONE, 14(4), e0215095. https://doi.org/10.1371/journal.pone.0215095

Samani, M. M., Agboada, D., Jamil, A., Kuo, M. F., & Nitsche, M. A. (2019). Titrating the neuroplastic effects of cathodal transcranial direct current stimulation (tDCS) over the primary motor cortex. Cortex, 119, 350–361.

Santarnecchi, E., Brem, A. K., Levenbaum, E., Thompson, T., Kadosh, R. C., & Pascual-Leone, A. (2015). Enhancing cognition using transcranial electrical stimulation. Current Opinion in Behavioral Sciences, 4, 127–178. https://doi.org/10.1016/j.cobeha.2015.06.003

Schroeder, P. A., Dresler, T., Bahnmueller, J., Artemenko, C., Cohen Kadosh, R., & Nuerk, H.-C. (2017). Cognitive Enhancement of Numerical and Arithmetic Capabilities: a Mini-Review of Available Transcranial Electric Stimulation Studies. Journal of Cognitive Enhancement, 1(1), 39–47. https://doi.org/10.1007/s41465-016-0006-z

Simonsmeier, B. A., Grabner, R. H., Hein, J., Krenz, U., & Schneider, M. (2018). Electrical brain stimulation (tES) improves learning more than performance: A meta-analysis. Neuroscience and Biobehavioral Reviews, 84(November 2017), 171–181. https://doi.org/10.1016/j.neubiorev.2017.11.001

Snowball, A., Tachtsidis, I., Popescu, T., Thompson, J., Delazer, M., Zamarian, L., … & Kadosh, R. C. (2013). Long-term enhancement of brain function and cognition using cognitive training and brain stimulation. Current Biology, 23(11), 987–992.

Turi, Z., Ambrus, G. G., Ho, K. A., Sengupta, T., Paulus, W., & Antal, A. (2014). When size matters: large electrodes induce greater stimulation-related cutaneous discomfort than smaller electrodes at equivalent current density. Brain stimulation, 7(3), 460–467.

Turi, Z., Boayue, N. M., Aslaksen, P., Antal, A., Paulus, W., Groot, J., Hawkins, G. E., Forstmann, B., Opitz, A., & Thielscher, A. (2018). Supplemental Material : Blinding is compromised for transcranial direct current stimulation at 1 mA for 20 minutes in young healthy adults. 1–7.

Turi, Z., Csifcsák, G., Boayue, N. M., Aslaksen, P., Antal, A., Paulus, W., Groot, J., Hawkins, G. E., Forstmann, B., Opitz, A., Thielscher, A., & Mittner, M. (2019). Blinding is compromised for transcranial direct current stimulation at 1mA for 20 min in young healthy adults. European Journal of Neuroscience, 50(8), 3261–3268. https://doi.org/10.1111/ejn.14403

Wagenmakers, E.-J., Marsman, M., Jamil, T., Ly, A., Verhagen, J., Love, J., Selker, R., Gronau, Q. F., Smíra, M., Epskamp, S., Matzke, D., Rouder, J. N., & Morey, R. D. (2018). Bayesian inference for psychology. Part I: Theoretical advantages and practical ramifications. Psychon Bull Rev, 25, 35–57. https://doi.org/10.3758/s13423-017-1343-3

Wallace, D., Cooper, N. R., Paulmann, S., Fitzgerald, P. B., & Russo, R. (2016). Perceived comfort and blinding efficacy in randomised sham-controlled transcranial direct current stimulation (tDCS) trials at 2 mA in young and older healthy adults. PLoS ONE, 11(2), 1–16. https://doi.org/10.1371/journal.pone.0149703

Zeng, F. G., Tran, P., Richardson, M., Sun, S., & Xu, Y. (2019). Human sensation of transcranial electric stimulation. Scientific reports, 9(1), 1–12.

