## Supplementary materials for "Blinding efficacy and adverse events following repeated transcranial alternating current, direct current, and random noise stimulation"

*Table 1. The AEs that did not demonstrate enough occurrences within each stimulation condition to warrant analysis. A black cell indicates that there were enough instances of an AE for a stimulation type within an experiment for analysis..*

| | **Experiment 1** | | **Experiment 2** | | |
|---|---|---|---|---|---|
| | **tDCS** | **tRNS** | **tDCS** | **tRNS** | **tACS** |
| Pain (24hr) | ■ | ■ | ■ | ■ | ■ |
| Pain (post) | ■ | ■ | ■ | | ■ |
| Irritation (24hr) | ■ | ■ | ■ | ■ | |
| Irritation (post) | ■ | ■ | ■ | ■ | |
| Concentration (24hr) | ■ | ■ | | | ■ |
| Concentration (post) | | | | | ■ |
| Nervousness | ■ | ■ | ■ | ■ | ■ |
| Nausea | ■ | ■ | ■ | ■ | ■ |
| Else | ■ | ■ | ■ | ■ | |
| Other | ■ | ■ | ■ | ■ | ■ |

*Table 2. Total number of AE incidence across all sessions for each tDCS stimulation group (active and sham) at each site during Experiment 1.*

| Site | Stimulation | Pain (24hr) | Pain (post) | Irritation (24hr) | Irritation (post) | Concentration (24hr) | Concentration (post) |
|---|---|---|---|---|---|---|---|
| Harvard | tDCS | 0 | 1 | 0 | 0 | 0 | 0 |
| Harvard | Sham | 0 | 1 | 0 | 0 | 0 | 0 |
| Honeywell | tDCS | 3 | 11 | 3 | 15 | 3 | 18 |
| Honeywell | Sham | 0 | 1 | 0 | 8 | 4 | 4 |
| NEU | tDCS | 1 | 8 | 0 | 5 | 0 | 32 |
| NEU | Sham | 0 | 0 | 1 | 0 | 0 | 30 |
| Oxford | tDCS | 4 | 19 | 0 | 1 | 3 | 6 |
| Oxford | Sham | 0 | 4 | 0 | 0 | 0 | 0 |
| Total | | 0.66% | 3.69% | 0.33% | 2.38% | 0.82% | 7.39% |

| Site | Stimulation | Sensation | Nervousness | Nausea | Else | Other |
|---|---|---|---|---|---|---|
| Harvard | tDCS | 3 | 0 | 0 | 3 | 1 |
| Harvard | Sham | 2 | 0 | 0 | 1 | 0 |
| Honeywell | tDCS | 44 | 4 | 0 | 6 | 1 |
| Honeywell | Sham | 10 | 0 | 4 | 4 | 0 |
| NEU | tDCS | 204 | 3 | 3 | 19 | 1 |
| NEU | Sham | 97 | 2 | 0 | 7 | 0 |
| Oxford | tDCS | 20 | 0 | 0 | 0 | 2 |
| Oxford | Sham | 15 | 0 | 0 | 0 | 1 |
| Total | | 32.43% | 0.74% | 0.57% | 3.28% | 0.49% |

*Table 3. Total number of AE incidence across all sessions for each tDCS stimulation group (active and sham) at each site during Experiment 2.*

| Site | Stimulation | Pain (24hr) | Pain (post) | Irritation (24hr) | Irritation (post) | Concentration (24hr) | Concentration (post) |
|---|---|---|---|---|---|---|---|
| Harvard | tDCS | 3 | 5 | 0 | 2 | 14 | 10 |
| Harvard | Sham | 0 | 13 | 0 | 8 | 35 | 20 |

| | | | | | | | |
|---|---|---|---|---|---|---|---|
| Honeywell | tDCS | 0 | 0 | 0 | 0 | 2 | 6 |
| Honeywell | Sham | 3 | 4 | 0 | 3 | 6 | 8 |
| NEU | tDCS | 1 | 8 | 0 | 2 | 37 | 24 |
| NEU | Sham | 1 | 13 | 4 | 1 | 51 | 34 |
| Oxford | tDCS | 0 | 2 | 0 | 2 | 7 | 15 |
| Oxford | Sham | 0 | 6 | 0 | 6 | 18 | 21 |
| Total | | 0.58% | 3.68% | 0.29% | 1.73% | 12.27% | 9.96% |

| Site | Stimulation | Sensation | Nervousness | Nausea | Else | Other |
|---|---|---|---|---|---|---|
| Harvard | tDCS | 37 | 0 | 0 | 3 | 0 |
| Harvard | Sham | 107 | 0 | 0 | 1 | 0 |
| Honeywell | tDCS | 17 | 0 | 0 | 8 | 0 |
| Honeywell | Sham | 25 | 0 | 0 | 9 | 0 |
| NEU | tDCS | 20 | 1 | 0 | 3 | 0 |
| NEU | Sham | 49 | 0 | 0 | 8 | 0 |
| Oxford | tDCS | 1 | 0 | 0 | 3 | 0 |
| Oxford | Sham | 11 | 0 | 2 | 9 | 0 |
| Total | | 19.26% | 0.07% | 0.14% | 3.17% | 0.00% |

*Table 4. Total number of AE incidence across all sessions for each tRNS stimulation group (active and sham) at each site during Experiment 1.*

| Site | Stimulation | Pain (24hr) | Pain (post) | Irritation (24hr) | Irritation (post) | Concentration (24hr) | Concentration (post) |
|---|---|---|---|---|---|---|---|
| Harvard | tRNS | 0 | 0 | 0 | 0 | 0 | 0 |
| Harvard | Sham | 0 | 0 | 0 | 0 | 0 | 0 |
| Honeywell | tRNS | 3 | 17 | 1 | 4 | 0 | 16 |
| Honeywell | Sham | 2 | 6 | 0 | 4 | 0 | 7 |
| NEU | tRNS | 1 | 18 | 0 | 5 | 0 | 43 |
| NEU | Sham | 0 | 3 | 1 | 1 | 0 | 14 |
| Oxford | tRNS | 0 | 0 | 0 | 0 | 0 | 0 |

| | | | | | | | |
|---|---|---|---|---|---|---|---|
| Oxford | Sham | 1 | 0 | 0 | 0 | 0 | 0 |
| Total | | 0.55% | 3.44% | 0.16% | 1.09% | 0.00% | 6.25% |

| Site | Stimulation | Sensation | Nervousness | Nausea | Else | Other |
|---|---|---|---|---|---|---|
| Harvard | tRNS | 2 | 0 | 0 | 2 | 1 |
| Harvard | Sham | 1 | 0 | 0 | 1 | 0 |
| Honeywell | tRNS | 31 | 1 | 2 | 14 | 1 |
| Honeywell | Sham | 17 | 1 | 0 | 2 | 2 |
| NEU | tRNS | 199 | 7 | 5 | 27 | 0 |
| NEU | Sham | 78 | 4 | 0 | 2 | 0 |
| Oxford | tRNS | 0 | 0 | 0 | 0 | 1 |
| Oxford | Sham | 0 | 0 | 0 | 0 | 0 |
| Total | | 25.63% | 1.02% | 0.55% | 3.75% | 0.39% |

*Table 5. Total number of AE incidence across all sessions for each tRNS stimulation group (active and sham) at each site during Experiment 2.*

| Site | Stimulation | Pain (24hr) | Pain (post) | Irritation (24hr) | Irritation (post) | Concentration (24hr) | Concentration (post) |
|---|---|---|---|---|---|---|---|
| Harvard | tRNS | 1 | 3 | 0 | 1 | 34 | 29 |
| Harvard | Sham | 0 | 5 | 0 | 1 | 27 | 19 |
| Honeywell | tRNS | 0 | 2 | 0 | 2 | 1 | 2 |
| Honeywell | Sham | 3 | 6 | 0 | 3 | 4 | 9 |
| NEU | tRNS | 2 | 12 | 0 | 4 | 27 | 25 |
| NEU | Sham | 4 | 11 | 2 | 4 | 85 | 55 |
| Oxford | tRNS | 0 | 6 | 1 | 0 | 9 | 21 |
| Oxford | Sham | 7 | 7 | 9 | 6 | 7 | 6 |
| Total | | 1.12% | 3.42% | 0.79% | 1.38% | 12.75% | 10.91% |

| Site | Stimulation | Sensation | Nervousness | Nausea | Else | Other |
|---|---|---|---|---|---|---|
| Harvard | tRNS | 19 | 0 | 0 | 1 | 0 |
| Harvard | Sham | 27 | 1 | 0 | 0 | 0 |

| | | | | | | |
|---|---|---|---|---|---|---|
| Honeywell | tRNS | 6 | 1 | 0 | 2 | 0 |
| Honeywell | Sham | 21 | 3 | 3 | 6 | 0 |
| NEU | tRNS | 15 | 1 | 4 | 8 | # |
| NEU | Sham | 39 | 4 | 3 | 9 | 0 |
| Oxford | tRNS | 13 | 0 | 0 | 6 | 0 |
| Oxford | Sham | 6 | 0 | 0 | 5 | 0 |
| Total | | 9.59% | 0.66% | 0.66% | 2.43% | 0.00% |

*Table 6. Total number of AE incidence across all sessions for each tACS stimulation group (active and sham) at each site during Experiment 2.*

| Site | Stimulation | Pain (24hr) | Pain (post) | Irritation (24hr) | Irritation (post) | Concentration (24hr) | Concentration (post) |
|---|---|---|---|---|---|---|---|
| Harvard | tACS | 0 | 0 | 3 | 5 | 0 | 0 |
| Harvard | Sham | 1 | 0 | 10 | 22 | 0 | 0 |
| Honeywell | tACS | 0 | 0 | 2 | 2 | 0 | 1 |
| Honeywell | Sham | 4 | 0 | 13 | 15 | 4 | 1 |
| NEU | tACS | 0 | 1 | 8 | 16 | 0 | 0 |
| NEU | Sham | 1 | 0 | 17 | 22 | 0 | 4 |
| Oxford | tACS | 0 | 0 | 11 | 7 | 5 | 6 |
| Oxford | Sham | 2 | 0 | 13 | 20 | 8 | 4 |
| Total | | 0.32% | 0.04% | 3.05% | 4.32% | 0.67% | 0.63% |

| Site | Stimulation | Sensation | Nervousness | Nausea | Else | Other |
|---|---|---|---|---|---|---|
| Harvard | tACS | 10 | 0 | 0 | 1 | 0 |
| Harvard | Sham | 52 | 0 | 1 | 1 | 0 |
| Honeywell | tACS | 2 | 0 | 1 | 1 | 0 |
| Honeywell | Sham | 12 | 0 | 0 | 21 | 0 |
| NEU | tACS | 28 | 0 | 0 | 7 | 0 |
| NEU | Sham | 55 | 0 | 3 | 24 | 0 |
| Oxford | tACS | 7 | 0 | 5 | 5 | 0 |

| | | | | | | |
|---|---|---|---|---|---|---|
| Oxford | Sham | 47 | 0 | 1 | 7 | 0 |
| Total | | 8.43% | 0.00% | 0.44% | 2.65% | 0.00% |

| Site | Experiment | No. Participants | Mean Age | Age S.D. | Age (min) | Age (max) | % Female | Years of Education | Education S.D. | Video Games Hours | Video Games S.D. |
|---|---|---|---|---|---|---|---|---|---|---|---|
| Harvard | 1 | 101 | 23.6 | 6.75 | 18 | 65 | 64.4 | 15.7 | 2.79 | 1.88 | 3.74 |
| Harvard | 2 | 134 | 24.9 | 8 | 18 | 58 | 41.8 | 16.4 | 2.6 | 2.95 | 4.91 |
| Honeywell | 1 | 110 | 24.7 | 7.58 | 18 | 65 | 39.1 | 16.3 | 2.49 | 4.64 | 8.19 |
| Honeywell | 2 | 178 | 22.8 | 7.17 | 18 | 65 | 42.1 | 15.1 | 2.52 | 3.9 | 6.86 |
| NEU | 1 | 133 | 23.9 | 4.64 | 18 | 56 | 25.6 | 16.6 | 2.9 | 4.79 | 7.08 |
| NEU | 2 | 193 | 23.8 | 2.33 | 18 | 36 | 27.5 | 17.2 | 1.75 | 5.11 | 6.59 |
| Oxford | 1 | 95 | 26.2 | 7.15 | 18 | 60 | 50.5 | 17.1 | 2.87 | 2.08 | 4.83 |
| Oxford | 2 | 165 | 23.3 | 5.02 | 18 | 47 | 63.6 | 16.4 | 2.7 | 1.7 | 3.6 |

*Table 7. Demographics for each site during each experiment.*

### Side effects questionnaire

| Subject Initials/Number: | Date: | Visit: |
|---|---|---|
| Condition Assignment: | | |

| Do you have any of the following symptoms? | | | | | | | |
|---|---|---|---|---|---|---|---|
| Symptoms | Yes/No<br>If "yes" to any symptoms, fill out severity and relationship | | Severity<br>1= Absent<br>2 = Mild<br>3= Moderate<br>4= Severe | | Relationship<br>1 = None<br>2 = Remote<br>3 = Possible<br>4 = Probable<br>5 = Definite | | Comments/Description |
| | Pre | Post | Pre | Post | Pre | Post | |
| Any changes since yesterday (or last visit)? | | | | | | | |
| Pain (i.e. headache, scalp pain, discomfort ) | | | | | | | |
| Scalp Irritation (redness, burn) | | | | | | | |
| **Only ask the following Questions Post-tES** | | | | | | | |
| Is anything feeling out of the ordinary? | | | | | | | |
| Trouble concentrating, focusing, or thinking | | | | | | | |
| If **YES**: Is it due to fatigue or sleepiness? ___________ | | | | | | | |
| Sensations under electrodes (tingling, itching, burning, etc) | | | | | | | |
| Visual Changes and/or Sensations | | | | | | | |
| If **YES**: Would you describe it as flashing lights or stars? __________ | | | | | | | |
| Nervousness | | | | | | | |
| Nausea | | | | | | | |
| Is there anything else that you would like to tell me? | | | | | | | |
| **Did the subject have any other adverse effect during or post-tES ?<br>Yes No if YES then write a summary of the event below** | | | | | | | |

Co-Investigator Signature Date

### **Blinding questionnaire**

1. a: How difficult did you find the test today, on a scale of 1-10?

1. b: How motivated were you to do your best on the test today, on a scale of 1-10?

1. c: How well do you think you performed on the test today, on a scale of 1-10?

1.d: How well do you think this test assesses the skills you have been practicing?

2. a: Each participant in this study received either real electrical brain stimulation or placebo stimulation while they played the training game. Do you think you were receiving real stimulation? NB: This question does not relate to the session you had today.

| | |
|---|---|
| | Yes |
| | No |

2. b: How confident are you about your answer?

| | |
|---|---|
| | Guess |
| | Possible |
| | Probable |
| | Definite |

2. c: Why do you think so?

3. a: Before the start of this study, did you think it was possible to improve cognitive abilities through training?

| | |
|---|---|
| | Yes |
| | No |

3. b: Has your opinion changed over the course of this study?

| | |
|---|---|
| | Yes |
| | No |

3. c: How much do you feel you improved in the training, relative to the maximum increase you think is possible? NB: The *training* refers to the game you played while you were wearing the headset.

3. d: To what extent do you think the *training* influenced your performance on the *tests* you did today, on a scale of 1-10?

4. Since the start of this study, did you notice any differences in your abilities to do daily tasks, like studying, solving problems, or remembering things?

| | |
|---|---|
| | Yes |
| | No |

For which tasks and to what extent did you notice differences?

### Blinding confidence

We additionally tested participants confidence in their response to the question about stimulation blinding. Analysis on participants' confidence in their responses revealed that for tDCS, tRNS, and tACS, the model containing the site by stimulation interaction explained the data best ($BF_{10}$=221, $BF_{10}$=109, and $BF_{10}$=4811, respectively; fig. S1).

#### tDCS

The interaction effect indicated that whilst sham participants reported similar levels of confidence across all sites, active stimulation participants were more likely to report increased confidence in their responses at HAR, with an approximately equivalent increase at NEU (b=0.42, 95%HDI=[-0.79, 1.53]) and OX(b=-0.03, 95%HDI=[-1.23, 1.13]). However, participants at the HON site who received active tDCS had much less confidence in their response (b=-1.00, 95%HDI=[-2.19, 0.14]). Post-hoc hypothesis tests revealed that participants in the active stimulation group at NEU were substantially more likely to report higher confidence in their answer than the sham participants (b=0.82, S.E.=0.39, 95%CI=[0.19,1.46]). This increase was less substantial for HAR (b=0.40, S.E.=0.46, [-0.34,1.16]) or OX (b=0.37, S.E.=0.39, [-0.28, 1.00]). In contrast, HON saw substantially lower confidence ratings in their active stimulation group compared to their sham (b=-0.60, S.E.=0.39, [-1.24, 0.02]).

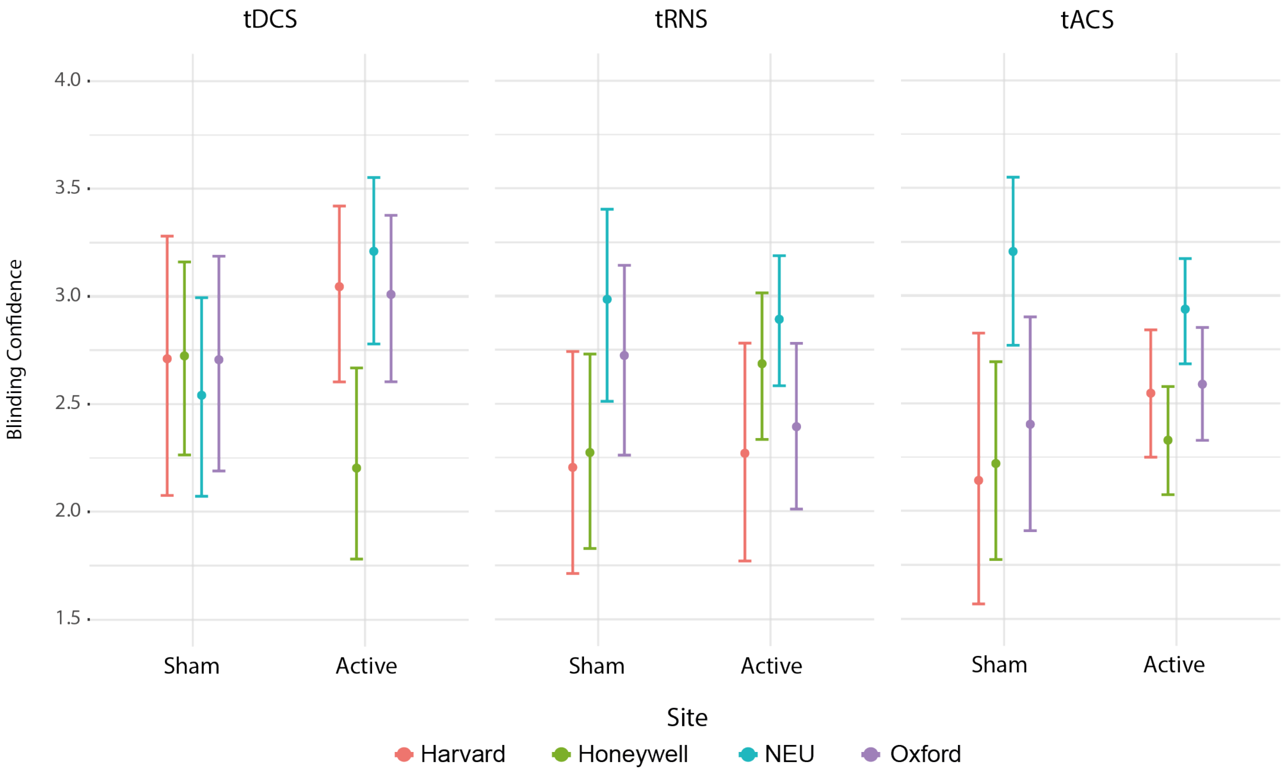

*Figure S1. Participants' confidence in their blinding response for each stimulation condition and site.*

**tRNS**

Sham participants in HAR and HON demonstrated equal levels of confidence in their response, whilst OX and NEU participants were more confident in their answers. In the stimulation group, confidence increased at the HON site (b=0.43, 95%HDI=[-0.71, 1.61]), but decreased at OX (b=-0.48, 95%HDI=[-1.64, 0.70]). There was little change in response at HAR or NEU (b=-0.20, 95%HDI=[-1.37, 0.96]). Post-hoc hypothesis tests revealed that stimulation groups differed most at HON (b=0.50, S.E.=0.36, [-0.09, 1.09]), followed by OX (b=-0.40, S.E.=0.38, [-1.02, 0.21]), with HON showing an increase in confidence amongst active participants and OX showing a decrease amongst active participants. By comparison, there was little difference between stimulation groups at NEU (b=-0.13, S.E.=0.37, [-0.73, 0.47]) or HAR (b=0.08, S.E.=0.47, [-0.70, 0.82]). Additionally, when compared against HAR, active participants at both NEU (b=0.77, S.E.=0.38, [0.38, 1.40]) and HON (b=0.51, S.E.=0.39, [-0.15, 1.14]) demonstrated substantially more confidence in their responses, whilst OX participants were approximately equivocal to HAR participants (b=0.15, S.E.=0.40, [-0.50, 0.81]).

Amongst participants who received tACS, participants in the active stimulation group were more confident than their sham counterparts at HAR, HON (b=-0.36, 95%HDI=[-1.48, 0.71]), and OX (b=-0.26, 95%HDI=[-1.36, 0.84]). However, confidence was lower amongst participants who received active tACS at NEU (b=-0.84, 95%HDI=[-1.91, 0.26]). Post hoc tests indicated that there were limited within-site differences between stimulation groups, with HAR (b=0.49, S.E.=0.44, [-0.25, 1.21]) and NEU (b=-0.35, S.E.=0.33, [-0.89, 0.19]) demonstrating larger changes than OX (b=0.22, S.E.=0.35, [-0.35, 0.79]) or HON (b=0.13, S.E.=0.33, [-0.42, 0.67]). Within the active stimulation group, NEU differed substantially from HAR (b=0.48, S.E.=0.24, [0.09, 0.88]) and HON (b=0.74, S.E.=0.22, [0.37, 1.11]). However, neither HON (b=-0.26, S.E.=0.24, [-0.66, 0.13]) nor OX (b=0.05, S.E.=0.24, [-0.35, 0.46]) differed from HAR.

#### Stimulation condition comparison

We ran a secondary analysis in which all stimulation conditions were included in the same model. Two factor predictors were created: tES, which had 3 levels (tRNS, tDCS, and tACS); and protocol, which had 2 levels (sham and real). Four models were run; a model containing only tES, a model containing only protocol, one with main effects for both, and one with main effects for both and the interaction effect. Each one was compared to a model that contained only an intercept. The model containing the main effects of both tES and protocol was the most favourable ($BF_{10}$ = 4420) compared to the null. Compared to tDCS, participants in the tRNS condition were less likely to report receiving active stimulation (b = -0.46, 95%HDI = [-0.79, -0.14]), where as those in the tACS condition were marginally more likely to report receiving active stimulation (b = 0.31, 95%HDI = [-0.05. 0.68]). Participants receiving sham stimulation were marginally less likely to report being in the active condition compared to participants receiving real stimulation (b = -0.31, 95%HDI = [-0.62, 0.01]).

**AE model out-of-sample results**

Model out-of-sample estimation ($elpd_{loo-ic}$) by using a leave-one-out cross validation approach calculated in the *brms* package in R. A model was considered favourable over the null if the difference in the elpd values (Δelpd) was larger than two times the difference in the standard errors of the elpd value (ΔS.E.). In situations where there were two models that fit this criterion (but did not reach the criterion of one model being a better fit than the other), the simpler model was selected.

**tDCS Experiment 1**

In Experiment 1, results indicated that model site was most likely for predicting skin irritation following tDCS (Δelpd=10.8, ΔS.E.=3.9). For changes in concentration following stimulation, the model containing site was most likely over the null (Δelpd=11.6, ΔS.E.=4.4). Finally, the model containing site was substantially more likely than the null for explaining any other AEs(Δelpd=6.1, ΔS.E.=2.5).

For all other AEs, models containing stimulation were not favourable over the null model or an alternate model not including stimulation.

**tDCS Experiment 2**

The model containing main effects for session and stimulation was favourable over the null (Δelpd=8.0, ΔS.E.=3.8). Similarly, the model containing session was more favourable for predicting post stimulation skin irritation data than the null model (Δelpd=10.5, ΔS.E.=3.9. Data indicated that a model with an interaction between stimulation and session was the most favourable over the null for predicting concentration in the 24 hours before the session (Δelpd=8.3, ΔS.E.=3.9). The model containing session was the best for predicting concentration AEs following stimulation (Δelpd=10.9, ΔS.E.=4.7) and sensation related AEs following stimulation (Δelpd=25.8, S.E.=5.6).

For all other AEs, the null model was favoured over any of the alternative models. Note, as there were no recorded "other" AEs, no analysis was run.

**tRNS Experiment 1**

The model containing the main effect of site was the most favourable over the null model for explaining the presence of pain AEs following stimulation

(Δelpd=6.0, ΔS.E.=1.9). Similarly, the model containing site was also favourable over the null for predicting concentration AEs after stimulation (Δelpd=15.2, ΔS.E.=3.5). The model containing site was the most favourable over the null in predicting sensation after stimulation (Δelpd=26.6, ΔS.E.=6.9). The model containing site was marginally more favourable over the null in predicting nervousness following tRNS stimulation (Δelpd=3.8, ΔS.E.=1.6). Finally, the model with the site main effect was better for predicting "else" AEs than the null model (Δelpd=4.2, ΔS.E.=1.9).

For all other AEs, the null model was favoured over the alternate models. As there were no recorded concentration AEs in the 24 hours prior to stimulation, no analysis was run.

**tRNS Experiment 2**

The model containing session was the best predictor of participants' pain during stimulation (Δelpd=17.9, ΔS.E.=5.8). This was also true for predicting participant concentration following stimulation where the session main-effect model was best compared to the null model (Δelpd=15.0, ΔS.E.=5.3. Participant reports of sensation also indicated that the model containing the main effect of session was favourable over the null model (Δelpd=15.7, ΔS.E.=6.0. In contrast to the other AEs reported for tRNS in Experiment 2, the model containing the main effects for stimulation and session was most favourable over the null (Δelpd=7.7, ΔS.E.=3.7).

For all other AE types, the null model was favoured over the alternate models. There were no recorded other AEs following stimulation, so no analysis was run.

**tACS**

Predicting irritation in the 24 hours preceding stimulation, the model containing session was the most likely (Δelpd=28.0, ΔS.E.=8.1). Concentration following stimulation was also best predicted by a model containing the main effect of session (Δelpd=17.4, ΔS.E.=5.6). Sensations in the period immediately following stimulation, the model containing the main effect of session was the best predictor (Δelpd=34.4, ΔS.E.=7.8). Similarly, nausea AEs following stimulation were best predicted by the session main effect only model (Δelpd=11.1, ΔS.E.=3.7). Finally, when asked about any "else" AEs experienced, once again, the session main effect only model was most likely (Δelpd=13.7, ΔS.E.=5.9).

For all other AE types, the null model was favoured over the alternate models.
